# Culture conditions antagonize lineage-promoting signaling in the mouse blastocyst

**DOI:** 10.1101/2020.03.31.016717

**Authors:** Tristan Frum, Amy Ralston

## Abstract

The mouse preimplantation embryo is a paradigm for discovery of the molecular principles governing formation of specific cell types during development. We show that conditions commonly used for *ex vivo* culture of preimplantation development are themselves antagonistic to a pathway that is critical for blastocyst lineage commitment.

## Introduction

Knowledge of the mechanisms that drive lineage decisions during mouse preimplantation development can significantly impact fields of stem cell and reproductive biology. Studies of the first three days of development have shown that the initially totipotent blastomeres of the mouse embryo adopt one of three cell fates by the blastocyst stage: trophectoderm (future placenta), primitive endoderm (future yolk sac endoderm), or epiblast (future fetus and additional extraembryonic tissues). Notably, the ultimate ratio of these three cell types is largely invariant among embryos, indicating that robust regulatory mechanisms exist. Remarkably, lineage specification is even achieved in incubator-grown preimplantation embryos, raised outside of the maternal environment, suggesting that the most critical regulatory mechanisms are intrinsic to the embryo. However, the possibility that culture conditions influence the embryo’s signaling environment cannot be formally excluded.

Major insight into the roles of cell signaling in preimplantation cell fate decisions has been provided by studies of the Fibroblast Growth Factor (FGF) signaling pathway (reviewed in Soszyńska *et al.*, 2019). Culturing embryos in FGF4 and Heparin (referred to as FGF4, hereafter) causes blastomeres to adopt primitive endoderm, over epiblast, fate (Yamanaka *et al.*, 2010). Importantly, the balance between primitive endoderm and epiblast fates is achieved through the exquisite regulation of the specific dose of FGF signaling experienced by the embryo as it develops (Krawchuk *et al.*, 2013).

The ability to culture preimplantation embryos in the presence of small molecule inhibitors or agonists of signaling has enabled discovery of the gene regulatory networks crucial for early mammalian development. For example, artificially tweaking the level of FGF/ERK signaling in embryos lacking specific transcription factors (*i.e.* transcription factor gene knockouts), can reveal whether transcription factors function upstream, downstream, or independently of FGF/ERK signaling. In addition, these assays have provided new insights into the developmental mechanisms in other mammals, including humans (Kuijk *et al.*, 2012; Roode *et al.*, 2012; Boroviak *et al.*, 2015; Piliszek *et al.*, 2017).

Culturing mouse embryos in agonists and antagonists of FGF/ERK signaling is routine, yet we note that this is typically performed in amino acid-containing KSOM medium containing 1 mg/mL bovine serum albumin (BSA). This raises the possibility that a bovine serum-derived impurity could alter the embryonic signaling environment. However, this possibility has not been explored.

## Results and Discussion

To better understand the embryonic signaling environment, we cultured embryos in the presence of 1 mg/mL polyvinyl alcohol (PVA), a synthetic macromolecule that can replace BSA in supporting blastocyst development (KSOM without BSA hereafter) (Biggers *et al.*, 1997; Jang *et al.*, 2007), and provide a more defined medium. In embryos cultured for 66 hours starting from the 2-cell stage (embryonic day E1.5) in KSOM without BSA, we observed quantitatively normal specification of trophectoderm, inner cell mass, epiblast and primitive endoderm cell types (Fig. 1A-C). This observation is consistent with prior evidence that PVA supports development of mouse blastocysts *in vitro* (Biggers et al., 1997).

**Figure 1.**
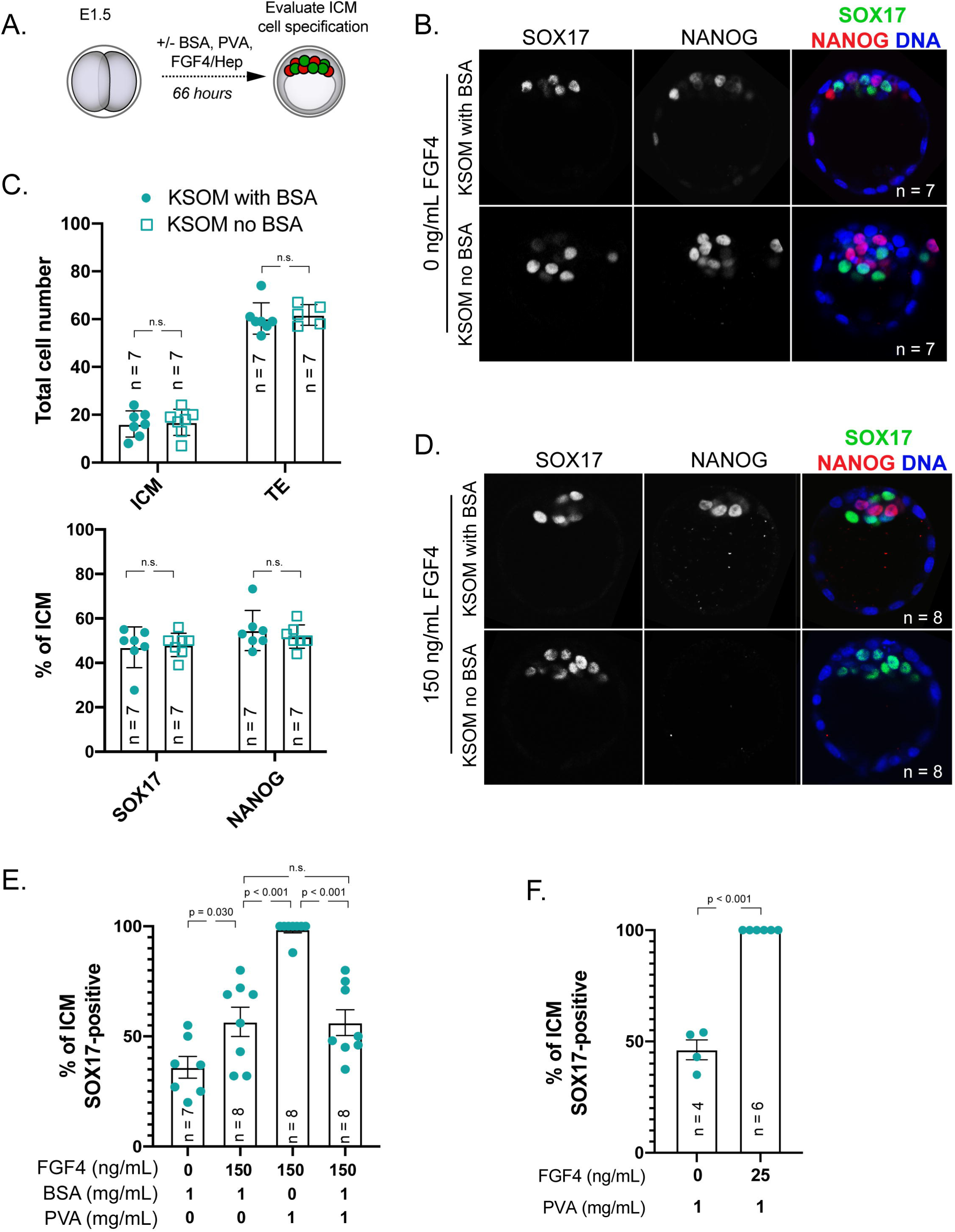
BSA antagonizes signaling by exogenous FGF4. A) Experimental design for embryos shown and analyzed in this figure. Red = NANOG indicative of epiblast cell fate, green = SOX17 indicative of primitive endoderm cell fate. B) Immunostaining for SOX17 to identify primitive endoderm (PE) cells and NANOG to identify epiblast (EPI) cells in embryos cultured for 66 hours starting from 2-cell stage in KSOM with 1 mg/mL BSA or without BSA (Millipore-Sigma MR-121-D and MR-107-D, respectively), but with 1 mg/mL added PVA (Sigma 360627). n = number of embryos examined. p = Student’s t-test. n.s. = not significant (p > 0.2). C) Quantification of the total inner cell mass (ICM) and trophectoderm (TE) cells in embryos cultured in indicated conditions and proportion of ICM cells contributing to PE and EPI in these embryos. Each symbol represents a single embryo, column = mean, and error bars = standard deviations. p = Student’s t-test. n.s. = not significant (p > 0.2). D) Immunostaining for SOX17 and NANOG in embryos cultured in KSOM + 150 ng/mL FGF4 (R&D Systems 235-F4-025) and 1 *µ*g/mL Heparin (Sigma H3149) in the presence or absence of 1 mg/mL BSA. Each symbol represents a single embryo, column = mean, and error bars = standard deviations. p = Student’s t-test. n.s. = not significant (p > 0.2). E) Quantification of lineage specification in embryos cultured in conditions indicated. Each symbol represents a single embryo, column = mean, and error bars = standard deviations. F) Response of embryos cultured in KSOM without 1 mg/mL BSA to low dose of exogenous FGF4 and 1 *µ*g/mL Heparin. Each symbol represents a single embryo, column = mean, and error bars = standard deviations. p = Student’s t-test.

Having established that KSOM without BSA supports blastocyst lineage specification, we then evaluated the sensitivity of blastocysts to exogenous FGF4 in the absence of BSA. Remarkably, embryos cultured in KSOM without BSA were sensitive to much lower doses of FGF4 (150 ng/mL) than embryos grown in the presence of BSA (Fig. 1D, E). Higher sensitivity to FGF4 is due to the absence of BSA, rather than the presence of PVA since BSA antagonized the effects of FGF4 signaling in the presence of PVA (Fig. 1E). Surprisingly, a dose as low as 25 ng/mL FGF4 was capable of driving epiblast cells to express primitive endoderm genes when embryos were cultured in KSOM without BSA (Fig. 1F), which is 20-40 times lower than the FGF4 dose typically used to convert epiblast to primitive endoderm. This observation suggests that higher doses of FGF4 are super-physiological. These observations strongly suggest that BSA is antagonistic to changes in embryonic cell fates induced by exogenous FGF4.

We do not yet understand the mechanism by which BSA interferes with signaling induced by exogenous FGF4. There is precedent that BSA can bind heparin, as well as extracellular signaling pathway members (Francis, 2010). We note that BSA does not appear to interfere with endogenous signaling, evidenced by the apparently normal lineage specification occurring in embryos cultured in KSOM with BSA. This could be because BSA in culture cannot penetrate the blastocyst to interfere with the highly localized FGF signaling occurring within the inner cell mass. We therefore caution that the routine supplementation with BSA could complicate studies aimed at understanding the roles and doses of signaling pathways in preimplantation development.

## Materials and Methods

### Mouse Stains

All animal care and husbandry for this study was performed in accordance with the guidelines established by the Institutional Animal Care and Use Committee at Michigan State University. Embryos in this study were obtained by timed natural matings of CD-1 mice (Charles River) which utilized males from 6 weeks to 6 months old, and females from 4 weeks to 3 months old.

### Embryo collection and culture

CD-1 embryos were collected at noon the day after observing the presence of a copulation plug (E1.5) by flushing oviducts with M2 medium (Millipore-Sigma, MR015D). Prior to transfer to the culture incubator, embryos were washed through 3x drops of embryo culture medium (KSOM with BSA (Millipore-Sigma, MR121D) or KSOM without BSA (Millipore-Sigma, MR107D) to remove M2 medium. KSOM without BSA was supplemented with 1 mg/mL PVA from a stock of 100 mg/mL PVA in water (Millipore-Sigma, P8136). Where indicated heparin (Millipore-Sigma, H3149) and FGF4 (R&D Systems, 235F4025) were added to embryo culture medium. Embryos were cultured under ES cell grade mineral oil (Millipore-Sigma, ES005C) for 66 hours at 37°C and 5% CO_2_. Embryo culture medium under mineral oil was equilibrated in the embryo culture incubator for at least 18 hours prior to the addition of embryos.

### Immunofluorescence and Confocal Microscopy

Embryos were fixed in 4% formaldehyde (Polysciences) for 10 minutes then immediately permeabilized for 30 minutes in 0.5% Triton X-100 (Millipore-Sigma, T8787) diluted in PBS. Embryos were stored in blocking buffer (10% FBS (Hyclone), 0.1% Triton X-100, PBS) for at least 18 hours and no longer than 5 days prior to incubation with primary antibodies. Primary antibodies were diluted in blocking buffer to the following concentrations: rabbit IgG anti-Nanog (Reprocell, RCAB002P) 1:400, goat IgG anti-SOX17 (R&D systems, AF1924) 1:2000 and applied overnight at 4°C. DyLight 488 (Jackson ImmunoResearch, 805485180) and Cyanine3 (Jackson ImmunoResearch, 711165152) conjugated antibodies targeting IgG were diluted at 1:400 in blocking buffer and incubated for 1 hour at ambient temperature to detect primary antibodies. Nuclei were labelled by 5-minute incubation with DRAQ5 (Cell Signaling, 4084) diluted 1:400 in blocking buffer. Confocal z-stacks spanning the entire embryo were collected with 5 *µ*m spacing on the Olympus FluoView FV1000 Confocal Laser Scanning Microscope system with 20x UPlanFLN objective (0.5 NA) and 5x digital zoom.

### Image Analysis

Image analysis was performed in ImageJ (imagej.net). Z-stacks from each embryo were arranged into a montage and then each cell was manually assigned a TE or ICM identity based on their position on the embryo surface (TE) or inside the embryo (ICM). Cells classified as ICM were further manually scored as SOX17 positive or NANOG positive. Pyknotic nuclei, indicating cell death, and mitotic nuclei were excluded from the analysis, and were less than 2% of all cells considered for analysis.

### Statistical Analysis

Unpaired two-tailed t-tests were performed in Graphpad Prism 8 for macOS (www.graphpad.com).

## Acknowledgements

We thank Dr. Jason Knott for insightful comments and suggestions on the manuscript. We apologize to authors whose work we were unable to cite due to space limitations.

## Author Contributions

AR conceived the study and wrote the paper. TF performed experiments, analyzed data, and produced the figure.

## Funding

This work was supported by the National Institutes of Health award R35 GM131759 to A.R.

## Declaration of Interests

The authors declare there is no conflict of interest that could be perceived as prejudicing the impartiality of the research reported.

